# Asynchronous suppression of visual cortex during absence seizures

**DOI:** 10.1101/282616

**Authors:** Jochen Meyer, Atul Maheshwari, Jeffrey Noebels, Stelios Smirnakis

## Abstract

Absence epilepsy is a common childhood disorder featuring frequent cortical spike-wave seizures with a loss of awareness and behavior. Using the calcium indicator GCaMP6 with *in vivo* 2-photon cellular microscopy and simultaneous electrocorticography, we examined the collective activity profiles of individual neurons and surrounding neuropil across all layers in V1 during spike-wave seizure activity over prolonged periods in *stargazer* mice. We show that most (∼80%) neurons in all cortical layers reduce their activity during seizures, whereas a smaller pool activates or remains neutral. Unexpectedly, ictal participation of identified single unit activity is not fixed, but fluctuates on a flexible time scale across seizures. Pairwise correlation analysis of calcium activity reveals a surprising lack of synchrony among neurons and neuropil patches in all layers during seizures. Our results demonstrate an asynchronous suppression of visual cortex during absence seizures, with major implications for understanding cortical network function during EEG states of reduced awareness.

---

Absence epilepsy interrupts normal cortical processing, producing reversible episodes of altered consciousness. Each seizure begins without warning, replacing planned motor movements with speech arrest and a vacant stare lasting only a few seconds, followed by sudden and complete recovery of awareness and intentional behavior. The events provide unique functional insight into the coupling of human perception and volition, and the cellular basis of this seizure type is steadily emerging from the study of genetic mouse models. The *stargazer* model, one of over 20 monogenic mouse mutants with this phenotype^1^, displays frequent, recurrent spike-wave seizures with behavioral arrest that are sensitive to blockade by ethosuximide^2^. Loss of the TARP subunit *Cacng2* in *stargazer* mice leads to mistrafficking of dendritic AMPA receptors in fast-spiking interneurons in the neocortex^3^ and thalamus^4,5^ and to remodeling of firing properties in thalamocortical circuitry that favor abnormal oscillations^6^. This loss of inhibition, particularly feedforward inhibition, is implicated in the pathophysiology of most monogenic models of absence epilepsy^1,7^. However, prior work in humans and animal models of absence epilepsy reveal inconsistent evidence of where and how cortical activity is modulated during seizures. While several studies in rats showed no activity changes in visual cortex^8^, or in somatosensory and motor cortex^9^, fMRI studies in humans have demonstrated increased ictal BOLD activity in the occipital cortex^10^, or biphasic activation and deactivation of large scale networks, including visual cortex^11,12^. Here, we combine simultaneous electrocorticography and 2-photon microscopy to overcome both the poor spatiotemporal resolution of fMRI and the limited number of neurons that can be simultaneously recorded with patch-clamp electrophysiology. Surprisingly, we found a suppression of both activity and synchrony in all layers of primary visual cortex of *stargazer* mice during electrographic spike-wave discharges.

Functional activity within large populations of GCaMP6 labeled neuron somata (n=29-132 per field) and patches of neuropil (n=4-18 per field) were readily visualized over prolonged periods in awake mice across all layers (Figure 1A, Supplementary Figure 1). Neurons with exceptionally low activity (∼12.6% of all labeled neurons) were excluded from the analysis (see methods). 22 datasets from 15 mice were analyzed (Supplementary Table), generating a total of 1366 distinct neurons and 222 patches of neuropil. For each dataset, the average activity of both neurons and neuropil in the ictal state (ΔF/F, mean±SEM of neurons = 5.3±1.7%; neuropil = 0.8±0.2%) was significantly lower than the activity in the interictal state (neurons = 8.1±2.2%; neuropil = 3.6±0.5%, paired t-test, p<0.001, Figure 2A, B). Calcium activity of individual neurons was then aligned to cortical EEG seizure onset or offset (Figure 1B), and distributions of calcium activity were compared between the interictal and ictal states using the Wilcoxon Rank Sum Test (see methods). Three patterns of ictal activity emerged (Figure 1A-D). In all cortical layers, most neurons had lower activity during ictal periods (mean±SEM, 82.3%±2.4%, “ictal-low” neurons in Figure 2C), while significantly fewer neurons showed either higher activity during ictal periods (“ictal-high”, 5.7%±0.9%, one-way ANOVA, p<0.005, Figure 2C) or no significant change (“neutral”, 12.1%±1.9%, p<0.005, Figure 2C). Neuronal firing rate was also estimated using a validated deconvolution algorithm^13,14^ (Supplementary Figure 2), revealing similar dominance of ictal-low over ictal-high neurons (Supplementary Figure 3). Deconvolution of calcium activity was validated with simultaneous cell-attached recordings and calcium imaging, showing excellent correlation between action potentials and visualized calcium transients (Supplementary Figure 2). Patch-clamp recordings from a subset of animals (n=16 putative pyramidal cells in 8 animals) further verified the predominant ictal-low firing patterns of L2/3 *stargazer* neurons (Ictal-low in 14/16, ictal-high in 2/16 (p=0.009, Wilcoxon paired rank-sum), Supplementary Figure 4). In addition, all neuropil patches, containing overlapping dendritic and axonal processes from many nearby neurons, were significantly ictal-low (Figure 2B). These findings demonstrate that the visual cortex is in a largely ‘suppressed state’ during absence seizure events.

**Figure 1:**
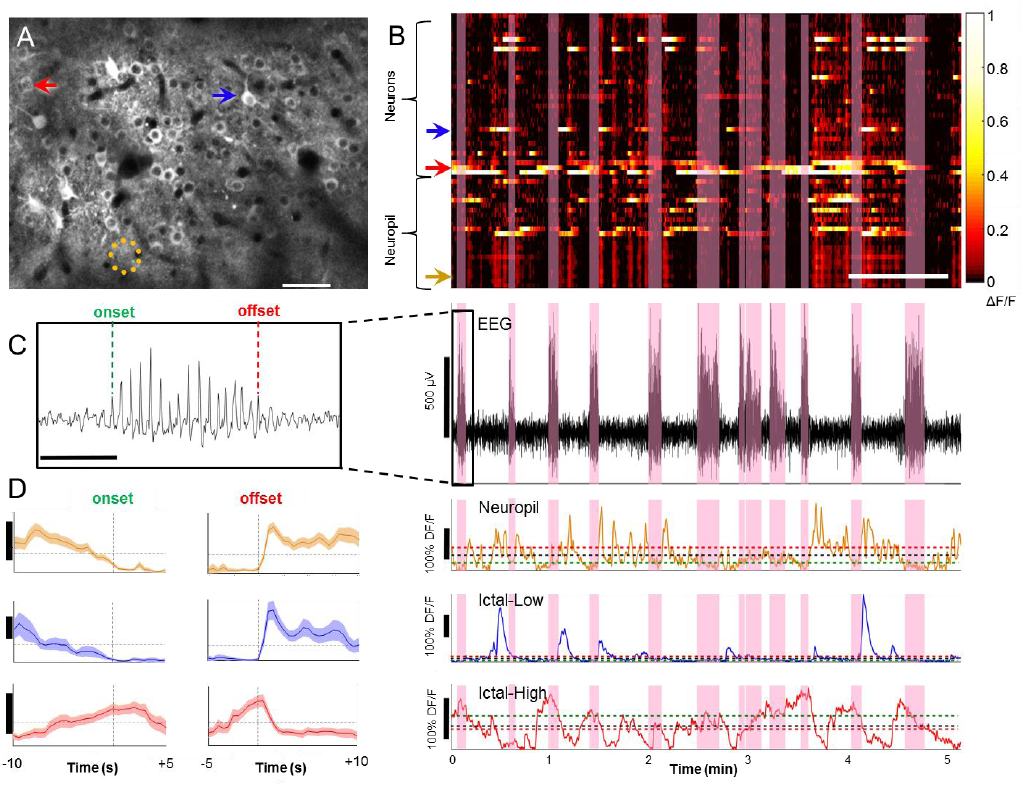
In vivo 2-photon microscopy and EEG in visual cortex of *stargazer* mice. **(A)** Typical field of view for *in vivo* imaging; GCaMP6-filled neurons in layer 2/3 of visual cortex (red arrow – neuron with high activity during seizures (ictal -high); blue arrow – neuron with low activity during seizures (ictal-low); orange dotted circle = neuropil patch. Bar = 50µm. **(B)** *<u>Top:</u>* Calcium activity for each neuron/neuropil patch over time with seizure epochs highlighted in pink (horizontal bar = 1 minute). *<u>Bottom:</u>* concomitant EEG, and traces of calcium activity from the depicted neuropil, ictal -low neuron, and ictal-high neuron shown in A. Mean ictal activity (green dashed line), mean interictal activity (red dashed line), and overall mean activity (black dashed line) are plotted. Inset **(C)** shows definition of seizure onset and offset at first and last spike of the seizure, respectively (bar = 1 second). **(D)** Average activity (mean±SEM, vertical bar = 20% ΔF/F) of an exemplary neuropil patch (top), ictal-low neuron (middle), and ictal-high neuron (bottom) are shown, time-locked to the seizure onset (left) and offset (right).

**Figure 2:**
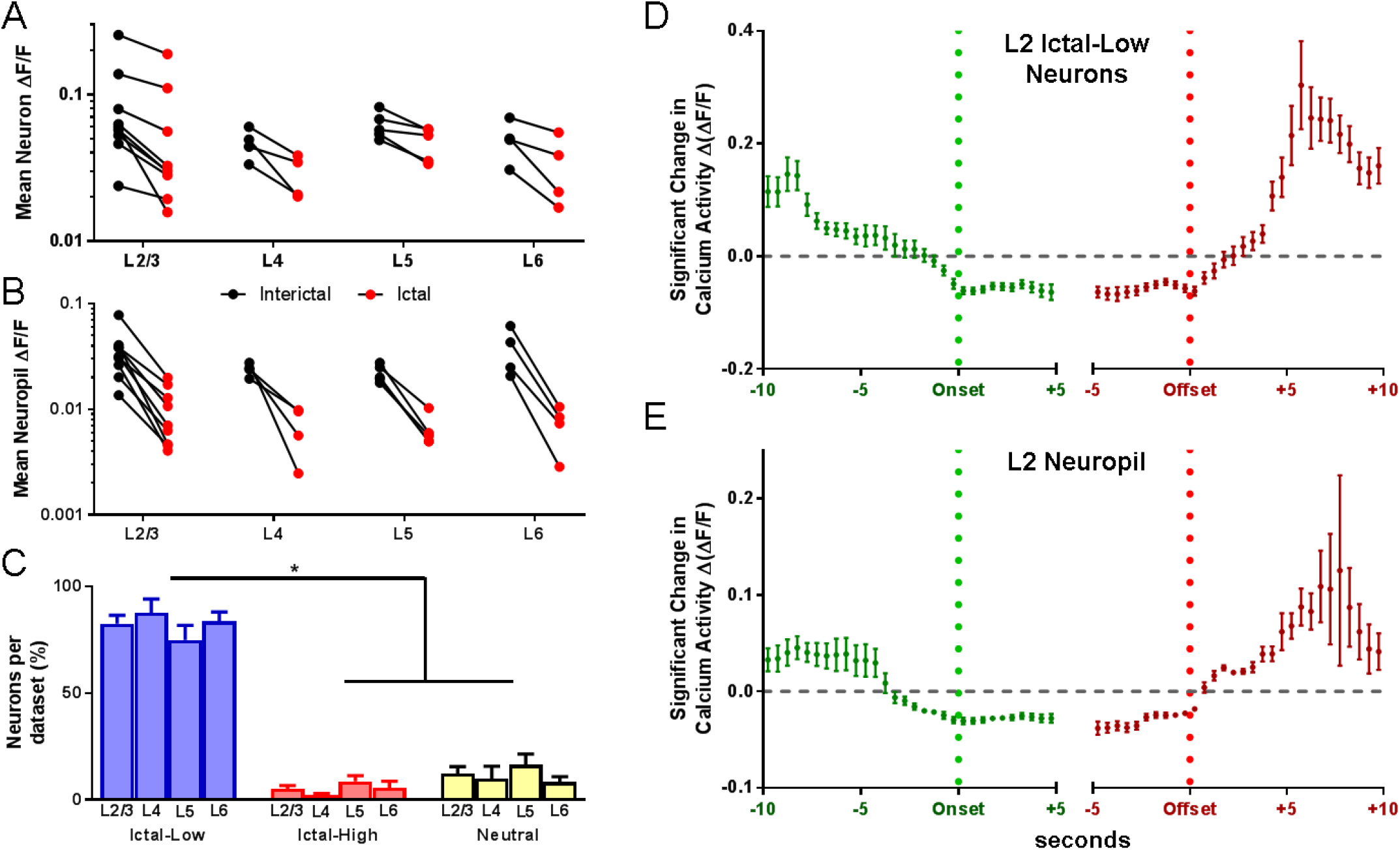
Activity changes in the ictal state. **(A)** Average calcium activity (ΔF/F) was significantly reduced in the ictal state compared to the interictal state for neurons across all layers. Line segments connect interictal (black disk) and ictal (red disk) mean activity within a field of view. Each comparison was statistically significant based on the Wilcoxon matched-pairs signed rank test (p<0.01) **(B)** Similar findings in neuropil (p<0.001 for all comparisons). **(C)** Significantly greater percentage of ictal-low compared to ictal-high or neutral neurons was seen across layers (p<0.0001 across datasets; 2-wayANOVA with Tukey’s multiple comparisons) **(D)** Temporal profile of significant change in ictal-low neuron activity compared to baseline, where Δ(ΔF/F) represents deviation from the average ΔF/F. Note that activity starts falling about 7 sec before seizure onset, returning to baseline following seizure offset (n/bin=9-155 neurons). **(E)** Temporal profile of change in neuropil activity shows that neuropil activity starts falling gradually more than 5 seconds prior to s ei zu re onset, with a similar return to baseline after seizure offset (n/bin=2-53 neuropil patches).

We next evaluated the time course of engagement for both neurons and neuropil by comparing calcium activity (ΔF/F) in half-second windows to a baseline (ΔF/F) generated from circularly shuffling the seizure time points (Kolmogorov-Smirnov test, p<0.05 after correction for multiple comparisons). Starting 10 seconds before and up to 5 seconds after seizure onset, an average of 33.1%±6.1% of neurons displayed significant change in activity (Δ(ΔF/F)) relative to seizure onset in at least one interval of two consecutive 0.5-sec bins, thereby indicating significant “participation” in seizure events. Using a similar 15-second window spanning 5 seconds prior to until 10 seconds following seizure offset, significant activity changes were found on average in 41.1%±6.7% of neurons. Layer 2 ictal-low neurons displayed gradually reduced activity starting about 7 seconds prior to seizure onset and then remained significantly below mean activity between 1 second prior to onset and 1 second after offset (Figure 2D). Neuropil activity displayed a gradual reduction earlier than neurons, dropping significantly below baseline 4 seconds prior to onset, but it returned to baseline around the same time as ictal-low neurons (Figure 2E). Neurons and neuropil in deeper layers showed similar reductions with Layers 4 and 5 dropping below baseline within several seconds of seizure onset, and Layer 6 at 2-3 seconds prior to onset (Supplementary Figure 5). In contrast to ictal-low neurons, only 21.7% of ictal-high and 31.6% of neutral neurons had significant activity in at least 2 consecutive 0.5-sec bins relative to baseline, with no consistent pattern relative to seizure onset and offset.

Given the variable participation of neurons relative to seizure onset and offset, we next asked to what degree each neuron remains within its overall classified activity profile over time, versus flipping to the opposite category. To do this, we compared ictal to interictal activity (ΔF/F) inside a sliding 7-minute window in datasets from all layers, which advanced from seizure to seizure over the course of the recording. The choice of a 7-minute window was empirically found to balance the power necessary for statistical analysis with temporal resolution. Figure 3A plots p-values for significance for each window in a typical ictal-high neuron, displaying 1 epoch of consecutive 7-minute windows which were in the predicted ictal-high state (red, 9 consecutive seizures). In this example, there was also 1 ictal-low epoch which stretched over 4 consecutive seizures (blue). As expected, both ictal-high and ictal-low neurons spent a significantly greater percentage of time in the predicted classification compared to the opposite classification (25% vs 6% over all neurons, p<0.0001, paired t-test, Figure 3B). Ictal-low neurons, however, had a significantly greater likelihood of remaining in the predicted classification (32% vs 9%, p=0.0124) and a significantly lower likelihood of being transiently classified in the opposite classification (0.5% vs 7%, p=0.00028) than ictal-high neurons across all layers (2-way ANOVA with post-hoc Bonferroni correction, Figure 3B). To evaluate the transient nature of seizure coupling over a longer time interval, the participation of defined Layer 2/3 neurons was evaluated for the same cortical window at multiple time points over the course of 8-12 days. Figure 3C-F presents an example. In total, for 3 chronically recorded animals with 3 recordings at least 24 hours apart, a single neuron had a 44.0±1.2% probability of changing its classification to any other class between any 2 time points. 60.6±2.9% of all neurons (ictal low, ictal high or neutral) changed at least once over the total time recorded. More than half of ictal-low neurons (57.9±2.9%) maintained a consistent classification throughout the 3 time points whereas ictal-high neurons were never found to be consistently ictal-high. Altogether, these data show that neocortical neuron activity can be loosely coupled to seizures over both short- and long-range time intervals, with ictal-low neurons having greater consistency than ictal-high neurons.

**Figure 3:**
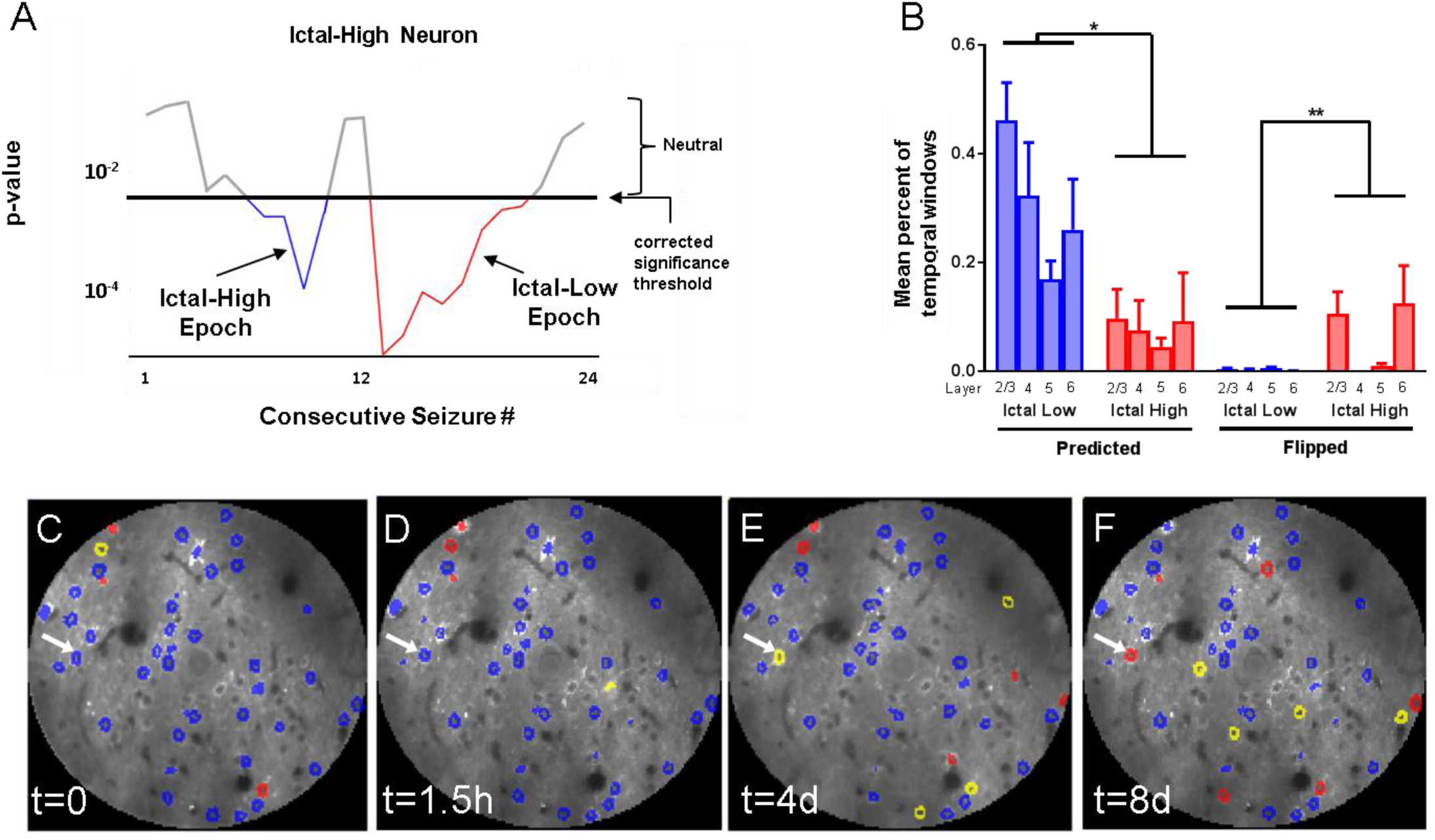
Temporal participation of neurons over minutes to days. **(A)** With a 7-minute moving window, a neuron with an overall classification of “ictal-high” had 1 significant ictal-high segment (red), periods of no significant difference (gray), and it also flipped to have 1 brief ictal-low epoch (blue). **(B)** Temporal windows were more often in the predicted than the flipped classification. In addition, ictal-low neurons had a smaller proportion of flipped classifications than ictal-high neurons (*p=0.028, **p<0.001). **(C)** Distribution of ictal-high and ictal-low neurons in one dataset, **(D)** 90 minutes later, **(E)** after 4 days, and **(F)** after 8 days in the same chronically imaged window. Note, for example, that one neuron starts as ictal low but then becomes neutral after 4 days and ictal high after 8 days (arrow). Blue=Ictal-low, Red=Ictal-High, Yellow=Neutral, Uncolored=Quiet (neurons with exceptionally low average activity in at least one of the 4 time points; see methods).

We next examined synchrony between and within nearby neurons and surrounding neuropil during interictal and ictal states using pairwise Pearson correlation coefficients. Since there is a mathematical reduction in correlation that occurs when average pairwise firing rates decrease, we corrected for baseline activity as previously described (see methods)^15,16^. The average correlation between pairs of neurons was significantly reduced in the ictal (0.09±0.01) compared to the interictal state (0.16±0.02), and neuropil to neuropil patch correlations were similarly reduced during seizures (ictal: 0.62±0.03, interictal: 0.81±0.02, Figure 4A-C, E, Wilcoxon matched-pairs signed rank test, p<0.005). As expected, intra-neuropil correlation coefficient strength was significantly greater than intra-neuron correlation in both the interictal and ictal states (p<0.001). When subdivided into specific groups, only ictal-low neurons were significantly less correlated in the ictal state, whereas ictal-high and neutral neurons had no significant change in correlation (Figure 4D). This pairwise reduction of correlation in ictal-low neurons had no relationship with the reduction in pairwise geometric mean calcium activity (r^2^=0.003, Supplementary Figure 6), further validating that the decrease in synchrony occurs independently of the reduced activity seen during the ictal state. Analysis with deconvolved traces (Supplementary Figure 6B, 7) yielded similar results. In addition, since locomotion itself has been associated with greater activity in the visual cortex^17^, it was important to demonstrate that our observations are not the indirect result of different locomotion rates in the interictal versus the absence (ictal) state. Two mice were analyzed after digitally subtracting image frames occurring during locomotion. As expected, only 0-1% of ictal frames coincided with locomotion, compared to 9.4% and 17.6% of frames in the interictal state of each dataset. Importantly, removal of locomotion-associated frames did not affect our basic observation that significantly lower neuronal activity and lower inter-neuronal synchrony occur in the ictal state (Supplementary Figure 8).

**Figure 4:**
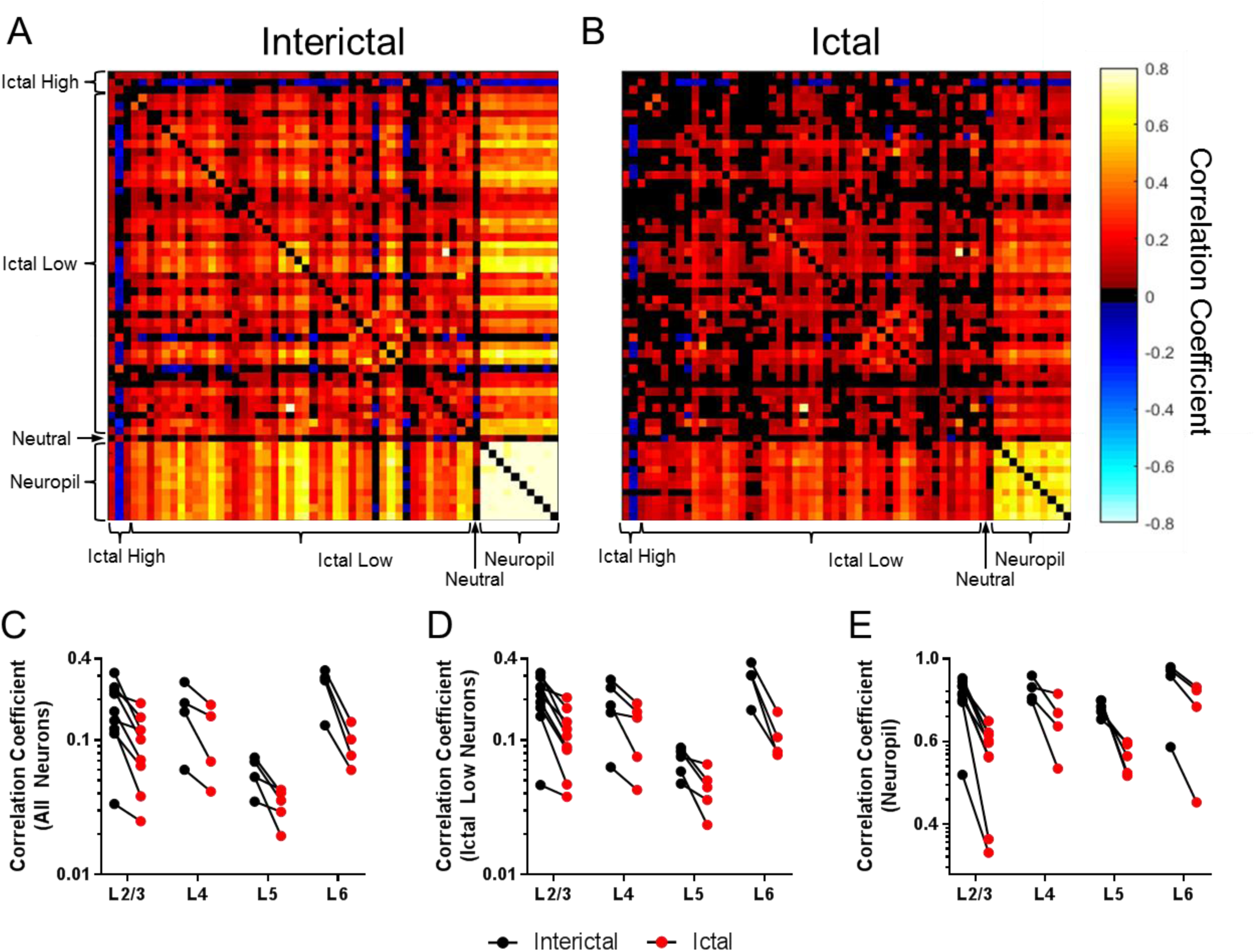
Pairwise synchrony is reduced in the ictal state. Pairwise correlation coefficients from a representative example of L2/3 neurons and neuropil in the **(A)** interictal and **(B)** ictal states. Across all datasets in all layers, there was a significant reduction in synchrony in the ictal state for **(C)** all neurons, **(D)** ictal low neurons, and **(E)** neuropil (*p*<0.005, Wilcoxon matched-pairs signed rank test).

Given the reduced synchrony between neurons during seizures seen at the temporal resolution of two-photon imaging (200 ms bins), we next evaluated to what degree single unit spikes correlate with seizure spikes on the EEG, at a temporal resolution of 20 ms bins. Pooled distributions of the timing of all action potentials recorded in Layer 2/3 (n=13,131 APs from 16 neurons) relative to the EEG spike times (taken at time 0) revealed only 2 (12.5%) neurons demonstrating a peak clustered around the time of EEG spikes, while the remaining 14 (87.5%) neurons had no significant temporal relation to the EEG spike times (D’Agostino ∼ Pearson normality test, *p*<0.05, Supplementary Figure 9). The mean likelihood of a neuron spiking within ±20 msec around an EEG spike was only 17.9±3.9% (mean±SEM). Given the overall low neuron-EEG spike synchrony, these findings do not support elevated synchrony of visual cortical neurons during spike-wave seizures at a higher temporal resolution.

To test if ictal classification may to some degree depend on cell type, we drove tdTomato expression in Dlx5/6-Cre (Dlx+) and Somatostatin-Cre (Som+) *stargazer* mice. Crossing Parvalbumin-Cre (PV+) with *stargazer* mice was limited by syntenic expression on chromosome 15^18^. We therefore employed the CLARITY technique^19^ with post-hoc immunostaining to identify parvalbumin-expressing interneurons (Figure 5). Out of 88 identified GABAergic interneurons across Layers 2-5, 84 (95.5%) were ictal-low (55 of 57 Dlx+, 12 of 13 Som+, and 17 of 18 PV+ interneurons), while the remainder were neutral neurons. No ictal-high neurons were found among all GABAergic neurons. Therefore, our data indicate that ictal classification of medial ganglionic eminence (MGE)-derived interneurons does not differ significantly from neuronal classification overall.

**Figure 5:**
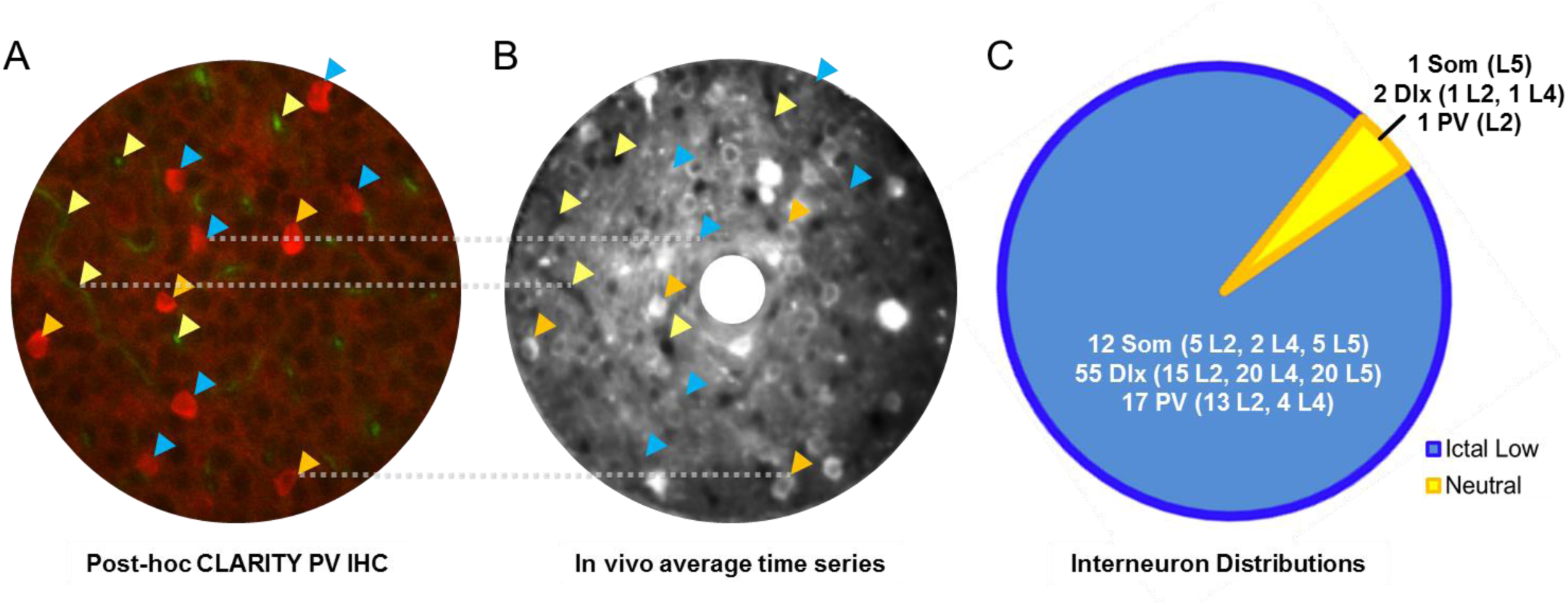
Interneurons identified *in vivo* and *ex vivo*. **(A)** Unperfused, clarified brain tissue allowed blood vessels to auto-fluoresce in the green channel (yellow arrowheads), while PV+ interneurons were immunostained and are shown in red (blue and orange arrowheads). **(B)** Co**-**registration with blood vessels (yellow arrowheads labeling both in-plane and perpendicular vessels) from an *in vivo* average of a time series revealed PV+ interneurons that were GCaMP6+ (orange arrowheads) versus GCaMP6-(blue). **(C)** Combined with interneurons identified *in vivo* due to expression of tdTomato under Somatostatin-Cre (Som) or Dlx-5/6-Cre (Dlx) drivers, the majority of identified interneurons were ictal low in nature.

Our results in the visual cortex differ from a previous electrophysiological description of neuronal firing in the somatosensory cortex^9^. In genetically inbred rats with absence epilepsy, intracellular recordings from a limited number (n=8-17) of single cortical neurons showed no significant change in firing rates during absence seizures^9,20^ with unit firing synchronized with the EEG spike discharge. Here, in contrast, simultaneous imaging of a large sample of V1 neurons revealed that mean neuronal activity (ΔF/F) is significantly reduced during the ictal state across all cortical layers. This is reflected in the fact that the large majority of neurons (81.5%) and neuropil are ictal-low, while ∼15% of neurons show no change in activity and only ∼5% increase their activity during seizures. Calcium signaling within neuropil has been shown to be more consistent with EEG activity^21^, and so our findings are consistent with the hypothesis that the electrographic spike-and-wave seizures of absence epilepsy are largely “inhibitory” seizures^22^. This is the first evidence at a cellular level that visual cortex is in a significantly hypoactive state during absence seizures, and is consistent with previous findings that cortical neurons have limited overall pathological excitation^23,24^ when they engage in absence compared to convulsive seizures^25–28^.

Previous fMRI studies of spike-wave epilepsy in humans have shown clear regional differences in activation, including signal increases during ictal activity in visual cortex^10^ or biphasic modulations in brain-wide networks, including the visual cortex^12^. One fMRI study in a rat model of absence epilepsy, however, showed no BOLD activity changes in visual cortex during seizures^8^. The variability among these reports and the difference from our results may be in part due to frequency dependent neurovascular coupling over a range of spike-wave discharge frequencies (3-11/sec) in these models^1,29^ and the lower spatio-temporal resolution of the BOLD signal. Our results also revealed that the cellular activity was loosely coupled to the seizures, with neurons dynamically changing their activity with respect to seizures over the course of minutes to days. This lack of a hard-wired cellular network correlate to a stereotyped EEG pattern is also consistent with recent findings in focal epilepsy in both mouse^28^ and human^30^ recordings.

Surprisingly, we also found a significant reduction in pairwise correlation strength during seizure events within the visual cortex, most pronounced in ictal-low neurons and surrounding neuropil, which was significant even after correcting for the reduced level of activity and reduced time spent in the ictal state. These findings are in direct contrast to spike-wave complexes associated with focal onset seizures, which show significantly increased pair-wise synchrony in the ictal state in vitro^31^ and in humans in vivo^26^. It is interesting to consider what this means in terms of information encoding^32,33^, and whether it could explain why absence seizures neither preclude the cortical transmission of evoked potentials^34^ nor have a post-ictal state. This issue will require further study in the context of stimulus presentation, taking also into account the effect that the accompanying subthreshold membrane voltage modulation is likely to have for information transmission across different areas.

Finally, it is of interest to speculate about the potential difference between ictal-low and ictal-high neurons. In a previous study of single units during focal seizures in patients with epilepsy, suppressed neurons were typically regular-spiking, presumably excitatory neurons, while increased activity was found in fast-spiking, presumably inhibitory neurons^35^. However, in our sampling of interneurons derived from the MGE (Dlx+, PV+, SOM+), the vast majority of these neurons had reduced activity in the ictal state. PV+ neurons in particular consistently showed reduced activity as predicted due to an AMPA receptor trafficking deficit previously described in these neurons^18^, which is consistent with a prior study showing that optogenetic inactivation of PV+ neurons in visual cortex leads to reduced pairwise correlation in cortical neurons^36^. Another source of suppression may be from a distinct class of interneurons including neurogliaform or vasoactive intestinal peptide (VIP)-expressing neurons. Since our findings were based on calcium imaging reflecting action potential firing, the apparent widespread synchrony on EEG may arise from synchronized subthreshold oscillations, which could be further dissected with simultaneous multi-cell patching and voltage-sensitive dyes. In line with the specific behavioral arrest associated with generalized spike-and-wave seizures, the asynchronously suppressed activity we observed in visual cortex provides a new framework for understanding cortical network malfunction leading to loss of awareness during absence ictal states, and for further exploring the cellular underpinnings of conscious experience.

## Acknowledgements

We acknowledge funding from: NINDS R21 NS088457 (JM and SS), NINDS K08 NS096029-01 (AM); and NINDS NS29709 (JLN).

## Data availability

Image and electrophysiology data will be made available by request to the authors.

## Supplementary Materials

### Materials and Methods

#### Surgical Implantation and chronic in vivo calcium imaging

All procedures were carried out according to an IACUC approved protocol. A minimum sample size of 8 mice was estimated using a power level set at 0.8 to detect an effect of at least 25% for each paired variable tested, with an estimated 4% variance between paired differences. AAV-1 GCaMP6M (11 animals) or GCaMP6S (1 animal) virus with a neuron-specific promoter (AAV1.Syn.GCaMP6m.WPRE.SV40, U Penn) was injected (100 nL) at 2 different depths (250 and 550 µm) using a Nanoject II automatic nanoliter injector through a burr-hole over the right V1 (2.5mm lateral and 1.5mm anterior to lambda) of 6-week old *stargazer* mice of either sex under isoflurane anesthesia. The deeper injection was located ∼300 µm further medial. This made it possible to image as deep as 760 µm below the dura due to the lack of overlying Gcamp6-expressing structures contributing out-of-focus fluorescence^37^. For some of the layer 4 through 6 imaging experiments, we used two different lines expressing Cre and tdTomato selectively in layer 4 (*Scnn1a*-Cre x Ai9) or layer 6 (*Ntsr1*-Cre x Ai9), and crossed these mice to the *stargazer* line. F1 offspring positive for Cre, Ai9 and heterozygous for the stargazin mutation were bred together, and a subset of their offspring were homozygous *stargazer* mutants expressing both Cre and tdTomato. 1 mm long, flat Ag/Ag-Cl electrodes were placed epidurally over the ipsilateral somatosensory cortex, 2 mm anterior to the middle of the craniotomy, and over the contralateral visual cortex; and a titanium headpost was permanently attached with dental cement. A reference electrode was implanted over the contralateral cerebellar hemisphere. A 3mm diameter glass cranial window was placed over the injected area and imaging was performed following a 4-6-week recovery period to allow sufficient cellular expression of the calcium indicator. Mice were imaged awake, head-posted in a holding frame and allowed to run freely on a circular treadmill. Raw calcium image series were acquired with a Prairie Ultima IV 2-photon microscope using a 25x objective, 1.1 NA, or a 16x objective, 0.8 NA, at 890 nm under spiral or raster galvo scanning mode (10 – 20 Hz frame rate). In 9 recordings from individual animals, we targeted layer 2/3 (mean depth below pia: 163 µm, range 100 – 240 µm). We also imaged 5 FOVs (from 4 animals) in layer 4 (mean depth below pia: 383 µm, range 360 – 395 µm), 5 FOVs (from 3 animals) in layer 5 (mean depth below pia: 570 µm, range 510 – 640 µm), and 4 FOVs (from 3 animals) in layer 6 (mean depth below pia: 730 µm, range 660 – 760 µm). Cells in layer 4 were identified either by using *stargazer* x *Scnn1a*-Cre/Ai9 line which expresses tdTomato in layer 4 pyramidal neurons, or by targeting the typical depth of layer 4 (330 – 480 µm)^38^, ensuring that cell bodies appeared relatively smaller than in layer 2/3. Layer 5 was identified either by using the *stargazer* x *Scnn1a*-Cre/Ai9 line and focusing right below layer 4, or by targeting the typical depth of layer 5 (480 – 680 µm)^38^, where pyramidal cell bodies are significantly larger than in layer 4. Layer 6 was identified either by using the *stargazer* x *Ntsr1*-Cre/Ai9 line which expresses tdTomato exclusively in layer 6 pyramidal neurons, or by targeting the typical depth of layer 6 (680 – 900 µm^31^), where the largest pyramidal cell bodies are smaller than in layer 5 (on average 214 µm^2^ in layer 5^39^ versus on average 133 µm^2^ in layer 6, based on our own measurements in two *stargazer* x *Ntsr1*-Cre mice). Laser output power under the objective was kept below 50mW, corresponding to ∼20% of power levels shown to induce lasting histological damage in awake mice^40^. In L5 and L6 FOVs, laser power was higher (up to 150 mW), but by visual inspection for signs of bleaching, cell swelling and other damage, we are confident that we did not cause physical harm to the neurons. This is because the power delivered per unit volume remains low as the total volume over which the power is distributed increases.

During imaging sessions, the EEG signal was sampled at 2 or 5 kHz with filtering cut-offs set at 1 Hz and 250 Hz. Seizures were detected by visual inspection by an experienced user (AM) in MATLAB (EEGLab) according to specific criteria (regular spike-wave burst structure, spike amplitude 1.5x baseline, spike frequency of 5-9 Hz, and a minimum duration of 0.5 seconds). The peaks of the first and last EEG spike were considered as the seizure onset and offset, respectively (Figure 1C).

In five animals, we recorded locomotion during the recording sessions using an angular velocity sensor. In addition, we monitored animal behavior by recording wide-field video of the mouse using an IR camera (Thorlabs DCC3240N) at 20 fps.

Patch-clamp recordings were performed in 8 *stargazer* and 2 control animals (n = 16 cells, 7 whole-cell, 9 cell-attached, plus 2 whole-cell (control)) for validation of the calcium-signal, as well for confirmation of the differences in ictal vs. interictal activity. We used borosilicate microelectrodes (4-7 MΩ) for current-clamp recordings and a previously published recipe for the current-clamp solution^15^, (in mM): 105 K-gluconate, 30 KCl, 10 HEPES, 10 phosphocreatine, 4 ATP-Mg, and 0.3 GTP. Data were acquired at 10 kHz and analyzed using custom-written MATLAB routines. Briefly, spike times were converted into firing rates and smoothed with a 100 msec sliding window. This window length was found empirically in patched neurons that expressed GCaMP6 by determining the highest correlation coefficient between their deconvolved traces and their actual smoothed spike rates using different smoothing window sizes. For further analysis, the resulting firing rate traces were subjected to the same algorithms used for analysis of the calcium activity traces described below.

#### Analysis of calcium imaging

The raw calcium fluorescence sequences were run through a custom x/y motion correction algorithm (MATLAB Inc.), based on a previously published method^41^. Briefly, motion parameters were estimated in the red channel in recordings from animals that had a subclass of interneurons labelled with tdTomato, otherwise the green channel was used, and motion corrected movies were visually inspected for errors and discarded if necessary. All image frames were registered to the average of the first 5 image frames using a sub-pixel registration method. Then, the correction offsets were applied to the green channel to reconstruct motion-corrected movies. Animals were resting on the treadmill during and between seizures. Since this seizure type is characterized by behavioral arrest, ictal movement was minimal. On average, only 7% of all frames per recording had some motion artifact that was corrected, with an average excursion of the frames exhibiting motion artifact of 1.5 µm. Gradual movement of <5 µm was reliably compensated for by the motion correction algorithm. Seizures with movement >5 µm occurring within 2 seconds before and after seizure onset or offset, were excluded from further analysis. To ensure that there was no significant contamination of cellular calcium activity by neuropil activity (especially in the z-axis), we tested a subset of recordings and confirmed that subtraction of neuropil activity from nearby cellular calcium signals did not affect the outcome of our analysis.

Regions of interest (ROIs: neuronal cell bodies and neuropil patches) were identified manually with ImageJ or semi-automatically using a custom MATLAB routine under supervision. The MATLAB routine was based on a previously published algorithm^42^: A seed point for a local ROI was placed manually with a circular ring to cover the cell body. Around the center of the cell, the algorithm then computes an intensity profile in polar coordinates. The final selection of the pixels for the ROI matches the maximal intensity along the polar profile. Neuropil patches were chosen to be larger than cell bodies because dendrites of local neurons labelled with GCaMP6, in contrast to other calcium indicators such as OGB, can have very high-amplitude fluorescence, which would dominate the signal of a neuropil patch if it consisted of relatively few pixels. Because we sought to analyze the signal comprised of contributions from a large number of axons and dendrites, comparable to LFP, neuropil patches for each dataset were, on average, 682±96 µm^2^ (mean±SEM). Neuropil patches were always further than 5 µm beyond the outline of ROIs defined as neurons.

Raw calcium traces for each ROI were created using the mean of all pixel intensities inside an ROI, and then high-pass filtered (0.1 Hz) and normalized by their individual baselines to ΔF/F values. To differentiate calcium transients from baseline fluctuations, the baseline (F) at time point t was defined as the mean of the bottom 10% of all data points within t±20sec, similar to prior work^42^. Noise was identified and removed from the ΔF/F signal by plotting a histogram of the negative data points of each ROI’s ΔF/F trace, fitting it with a half-Gaussian function, identifying the 0.5-SD distance below zero, and removing all data points below the peak of the Gaussian + 0.5-SD. We used simultaneous patch-clamp and calcium imaging data to calibrate this procedure. We also performed a similar analysis after deconvolving the calcium traces and found similar results. Firing rates were extrapolated from the ΔF/F traces using a modified version of 2 different deconvolution methods^13,14^, yielding consistent results. The method used to deconvolve the data we present in the text was based on a previously published method^13^ that uses 1) an iterative smoothing process to remove local low-amplitude peaks representing noise without distorting the ΔF/F signal stemming from calcium fluctuations, and 2) inverse filtering of the smoothed traces with an exponential kernel. The other method was used for further validation of the results and infers an approximation of the most likely spike train underlying the given fluorescence trace using an iterative expectation maximization algorithm.

We empirically determined a minimum level of activity for an ROI, below which it was not possible to determine whether the ROI had statistically significant firing events due to lack of data points, by calculating the sum of noise-corrected ΔF/F signal per minute for each ROI. ROIs with mean activity below this threshold (sum(ΔF/F)/min = 6 for GCaMP6M, sum(ΔF/F)/min = 22.5 for GCaMP6S) were deemed “quiet” (12.6±1.6% of all neurons) and excluded from further analysis^43^. The remaining ROIs were then classified as ictal-high, ictal-low or neutral by comparing all activity during the ictal state with all activity in the interictal state using the Wilcoxon rank-sum test (MATLAB) with Bonferroni correction for multiple comparisons. Comparison between and within group data was performed in Prism 5 (version 5.0d, GraphPad, CA, USA).

#### Temporal Analysis of calcium imaging

To minimize cross-contamination due to the relatively slow calcium signal dynamics, seizures (and adjacent inter-seizure intervals) that were shorter than 1.5 seconds, as well as seizures that were less than 6 seconds apart, were also excluded from analysis.

To determine the temporal evolution of each ROI’s classification around seizure onset and offset, calcium activity was aligned to seizure onset and offset separately, and null distributions of calcium activity were created by circularly shuffling the seizure time points 2,000 times while keeping the seizure length and calcium activity data constant (MATLAB). 30 bins of activity at 0.5-second intervals were created spanning from 10 seconds prior to until 5 seconds after seizure onset as well as from 5 seconds prior to until 10 seconds after seizure offset. The difference between the activity for each bin relative to seizure onset or seizure offset was compared against the corresponding bin of the null distribution to determine if the activity (ΔF/F) in a given ROI was significantly increased or decreased at that particular temporal window (MATLAB, Kolmogorov-Smirnov test, significance set at p<0.0017 (0.05/30) for correction of multiple comparisons). To be conservative, we undertook the further step of requiring that at least 2 neighboring 0.5-sec bins reached significance.

In order to characterize the behavior of each ROI on a finer temporal scale, we applied the same classification analysis to sliding windows of 7 minutes of activity along the duration of each recording. Each window was then advanced serially by one seizure. Seizure start and end times were then evaluated within any given 7-minute window, ictal time periods and interictal time periods were concatenated within each window and binned in 2-second intervals, and the Wilcoxon rank-sum test (MATLAB) was performed to compute significance between the ictal and interictal states.

#### Correlation analysis

Pair-wise Pearson correlation coefficients were calculated (MATLAB) separately for the interictal versus the ictal states as follows: The amplitude of each ROI’s ictal activity was normalized by the average amplitude of the ROI’s interictal activity. In order to correct for the mathematical reduced likelihood of correlation created by reduced activity, for each pair of neurons, we circularly shuffled one neuron’s concatenated ictal and interictal time points separately 2,000 times and computed null distributions of correlation coefficient strengths. The mean of the null distributions was then subtracted from the unshuffled original coefficients independently for the interictal and ictal states as previously described^16^. Alternatively, we randomly removed calcium-events from interictal (if interictal activity was higher) or ictal (if ictal activity was higher) time periods, until ictal and interictal average ΔF/F rates were less than 0.1% apart for each cell. Analyzing concatenated epochs versus averaging correlation coefficients from individual epochs yielded similar results, indicating that the concatenation process did not artificially affect correlations (data not shown).

#### Analysis of patch-clamp electrophysiological data

To determine whether patched neurons were more active during versus between seizures, we identified action potential (AP) times in voltage traces from cell-attached or whole-cell recordings (custom MATLAB routine) and calculated the ictal and interictal firing rates. APs were identified by searching for any time points t in the voltage trace that met the following criteria: i) V_m_(t+0.25 ms) > V_m_(t) + x, ii) mean(V_m_(t-pre_1_ to t-pre_2_)) < mean(V_m_(t+p_1_ to t+p_2_)) – α*x, and iii) mean(V_m_(t+post_1_ to t+post_2_)) < mean(V_m_(t-pre_1_ to t- pre_2_)) + β*x, where the threshold x was 38 mV, pre_1_ was 1.2 ms, pre_2_ was 0.3 ms, p_1_ was 0.25 ms, p_2_ was 0.35 ms, post_1_ was 2.5, post_2_ was 2.9 ms, α was 0.5 and β was 0.8 for whole-cell recordings, and x was 1.9 mV, pre_1_ was 0.8 ms, pre_2_ was 0.25 ms, p_1_ was 0.15 ms, p_2_ was 0.24 ms, post_1_ was 1.8, post_2_ was 2.1 ms, α was 0.55 and β was 0.45 for cell-attached recordings. In each recording, we identified the seizure spikes and used the spike times as zero-points to generate the peri-spike time histograms (from -100 ms to +100 ms) that indicate to what degree APs were time-locked to the spikes of a seizure.

## Code availability

Custom written MATLAB routines will be made available per request to the authors.

## Supplementary Materials: 1 table and 9 figures

**Supplementary Table.** Ictal/interictal characteristics and GCaMP6 indicator subtype used. ISI = inter-seizure interval

| Layer | Mouse | GCaMP6 type | Avg sz length (sec) | Avg ISI (sec) | # Sz | Total time (sec) | Total interictal (sec) | Total ictal (sec) |
| --- | --- | --- | --- | --- | --- | --- | --- | --- |
| 2/3 | 1 | slow | 2.9 | 33.1 | 46 | 1453 | 1319 | 133 |
|  | 2 | medium | 3.9 | 21.8 | 143 | 2381 | 1837 | 545 |
|  | 3 | medium | 7.0 | 32.1 | 47 | 1240 | 910 | 330 |
|  | 4 | medium | 4.5 | 36.0 | 32 | 1162 | 1017 | 145 |
|  | 5 | medium | 3.8 | 33.7 | 18 | 478 | 415 | 63 |
|  | 6 | medium | 3.1 | 58.4 | 25 | 1105 | 1028 | 76 |
|  | 7 | medium | 2.2 | 11.2 | 119 | 1587 | 1323 | 255 |
|  | 8 | medium | 3.5 | 11.0 | 95 | 1471 | 1036 | 332 |
|  | 9 | medium | 3.6 | 10.8 | 88 | 1249 | 940 | 309 |
| 4 | 6 | medium | 1.4 | 24.2 | 132 | 3434 | 3175 | 185 |
|  | 7 | medium | 2.7 | 31.6 | 40 | 1336 | 1232 | 104 |
|  | 9 | medium | 3.1 | 11.3 | 107 | 1533 | 1192 | 331 |
|  | 9 | medium | 2.7 | 13.5 | 53 | 1126 | 704 | 143 |
|  | 10 | medium | 3.8 | 16.6 | 75 | 1595 | 1227 | 280 |
| 5 | 7 | medium | 1.6 | 30.2 | 50 | 1575 | 1478 | 77 |
|  | 7 | medium | 2.0 | 7.7 | 157 | 1533 | 1202 | 313 |
|  | 9 | medium | 3.2 | 8.7 | 132 | 1460 | 1072 | 388 |
|  | 9 | medium | 3.4 | 8.1 | 135 | 1562 | 1080 | 459 |
|  | 10 | medium | 2.5 | 10.1 | 122 | 1548 | 1223 | 303 |
| 6 | 8 | medium | 2.9 | 15.2 | 96 | 1553 | 1261 | 211 |
|  | 8 | medium | 3.5 | 11.0 | 95 | 1471 | 1036 | 332 |
|  | 10 | medium | 3.3 | 8.8 | 129 | 1573 | 1131 | 424 |
|  | 11 | medium | 2.9 | 6.3 | 187 | 1737 | 1166 | 547 |
| **AVERAGE** |  |  | **3.2** | **19.6** | **92.3** | **1528.8** | **1217.6** | **273.3** |

**Supplementary Figure 1.**
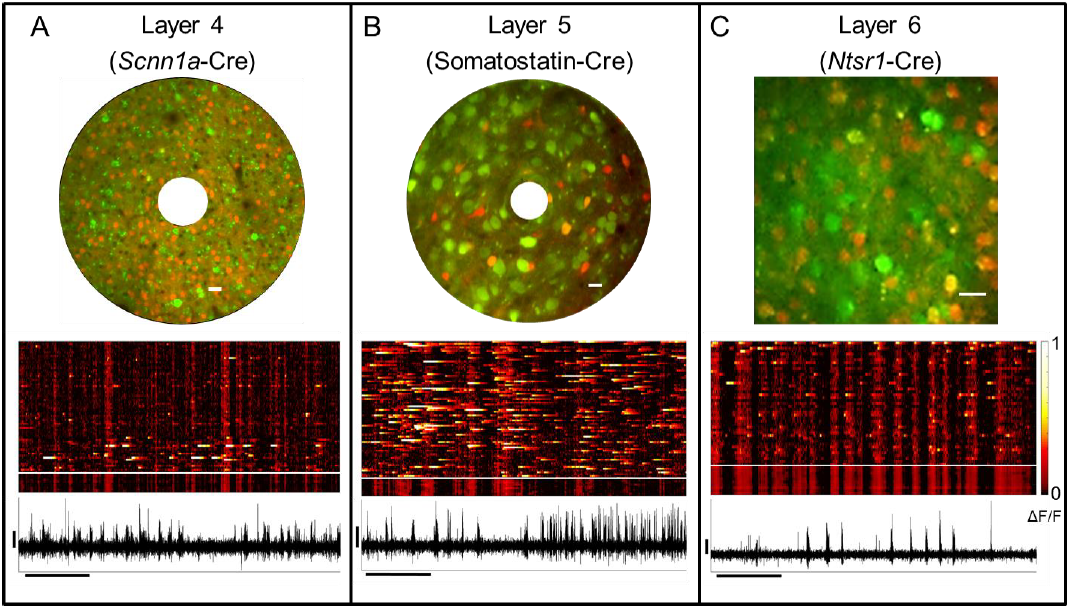
Deep layer imaging. Neurons and patches of neuropil were imaged in deeper layers using a combination of depth from the pial surface and layer-dependent expression of tdTomato. A typical field of view for each layer (above), raster plot of activity (middle, neurons above and neuropil below white line) and concomitant EEG (below) are seen in **(A)** Layer 4 (spiral scan), **(B)** Layer 5 (spiral scan), and **(C)** Layer 6 (line scan) of visual cortex (white scale bar = 20 µm, horizontal black scale bar = 10 seconds, vertical black scale bar = 200 µV).

**Supplementary Figure 2.**
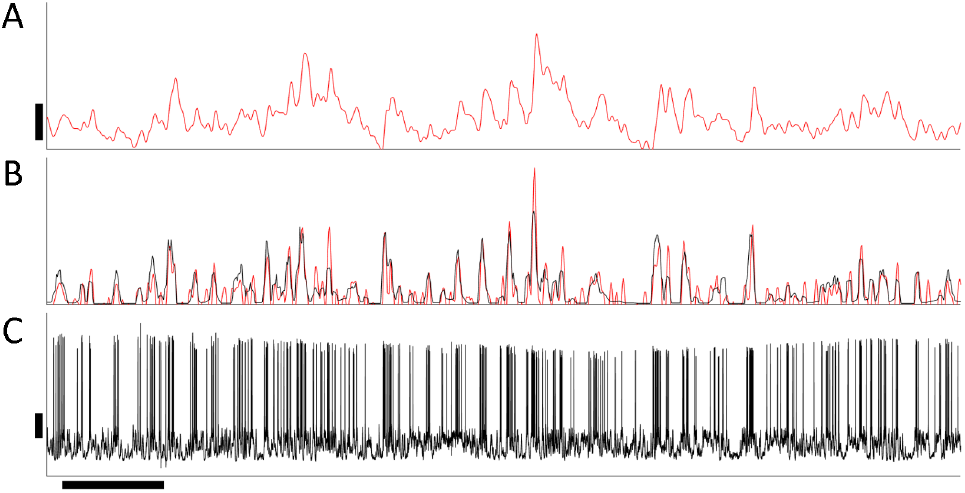
Correlation between action potentials and calcium activity in a *stargazer* mouse. A patched, GCaMP6m -filled neuron showing: **(A)** a representative ΔF/F trace (vertical scale bar = 20% ΔF/F), **(B)** the corresponding deconvolved activity trace (red), the extrapolated firing rate (black, A.U.), and **(C)** the corresponding membrane voltage trace (vertical scale bar: 20 mV). Horizontal scale bar = 10 sec. Correlation coefficients between deconvolved activity and extrapolated firing rates were ∼0.8 on average, and the percent of identified action potentials with deconvolved GCaMP6 activity were, on average, 80%, 97%, and 100% for singlets, doublets, and ≥triplets, respectively.

**Supplementary Figure 3.**
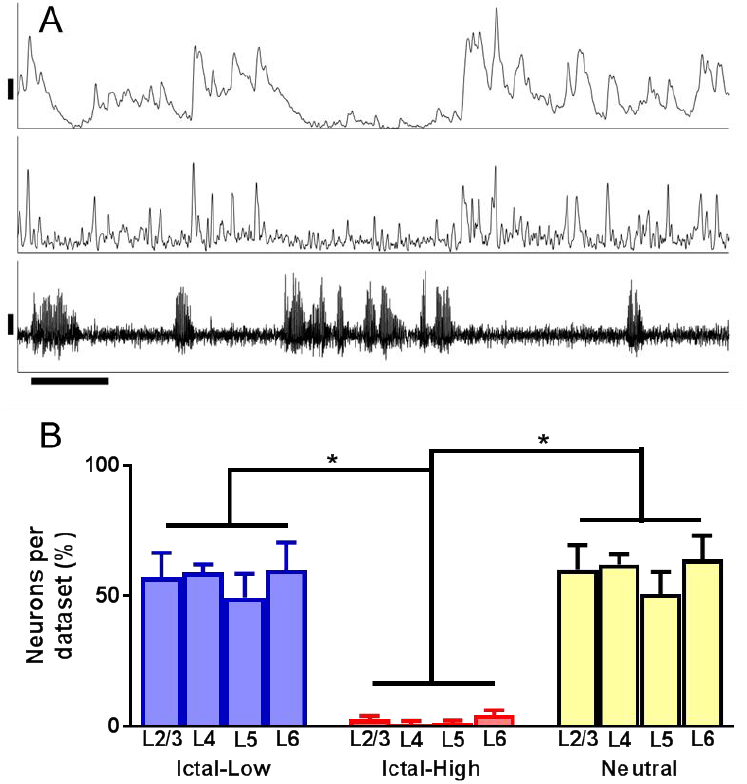
Deconvolution of the calcium signal yields similar results. **(A)** Example of activity traces for a single neuron before and after deconvolution (*top*, ΔF/F before deconvolution, vertical bar = 10%); *middle*: ΔF/F, deconvolved, A.U.; *bottom*: simultaneous EEG highlighting that lower activity during seizures can be seen both by analyzing the raw and deconvolved calcium recordings. Vertical bar: 200 µV, horizontal bar = 10 sec. **(B)** Significantly more ictal-low than ictal-high and neutral neurons (one-way ANOVA with post-hoc Bonferroni correction, *p<0.001) are observed after deconvolution. The lower percentage of ictal-low (56.1%) and ictal-high (2.3%) neurons is secondary to the fewer non-zero data points created as a byproduct of deconvolution. Note that in spite of this, the basic conclusions remain unchanged when using deconvolved data.

**Supplementary Figure 4.**
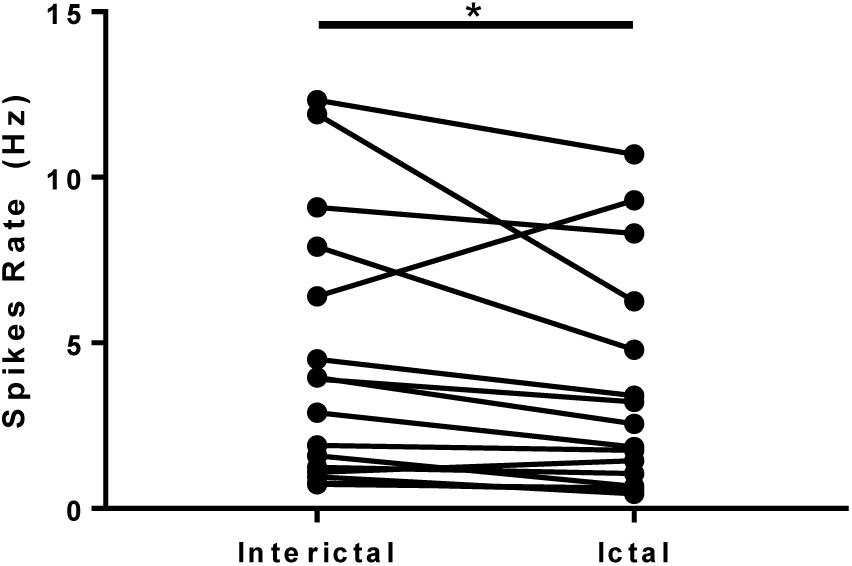
Patch-clamp recordings in a subset of animals corroborate the predominantly ictal-low character of L2/3 neurons. Mean ictal and interictal firing rates were calculated from single unit action potentials. 14 of 16 cells were deemed ictal low and 2 cells were ictal high; overall, the neurons were significantly suppressed (\**p*=0.0092, Wilcoxon matched pairs signed-rank test).

**Supplementary Figure 5.**
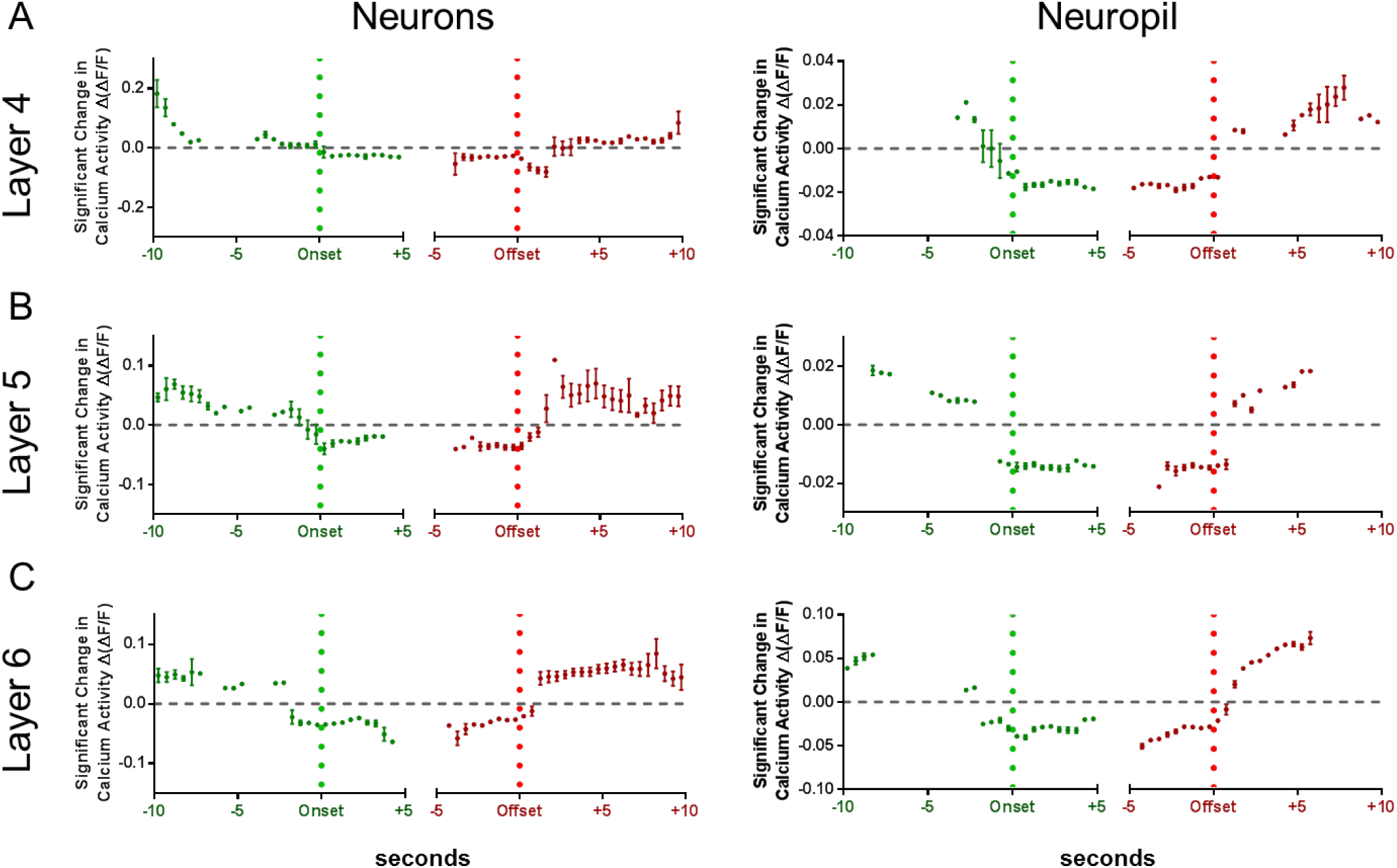
Temporal analysis of deep layers is similar to superficial layers. In **(A)** Layer 4, **(B)** Layer 5, and **(C)** Layer 6, ictal-low neurons and neuropil conthe ictal state, with reductions inthe ictal state, with reductions inthe ictal state, with reductions insistently reduce activity within a few seconds before onset and after offset. Layer 6 neurons cross below mean activity 2 seconds prior to onset, compared to Layer 4 neurons which cross below mean activity right at seizure onset.

**Supplementary Figure 6.**
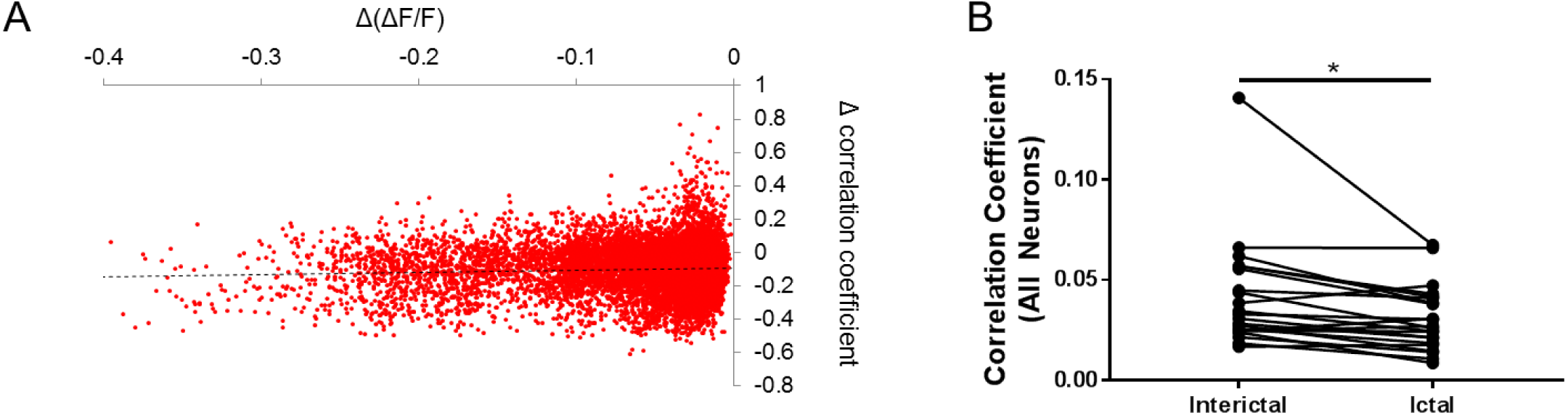
Validation of reduced synchrony in the ictal state. **(A)** There is no significant relationship between the geometric pairwise mean of reduced calcium activity and change in pairwise correlation coefficient (dotted line, r^2^=0.003), confirming that the observed decorrelation is not simply due to a change in firing rate. **(B)** Furthermore, after randomly removing activity until deconvolved traces show matching interictal and ictal rates for each neuron, the overall reduction in pairwise correlation in the ictal state remains (\**p*=0.0005, Wilcoxon matched pairs signed-rank test).

**Supplementary Figure 7.**
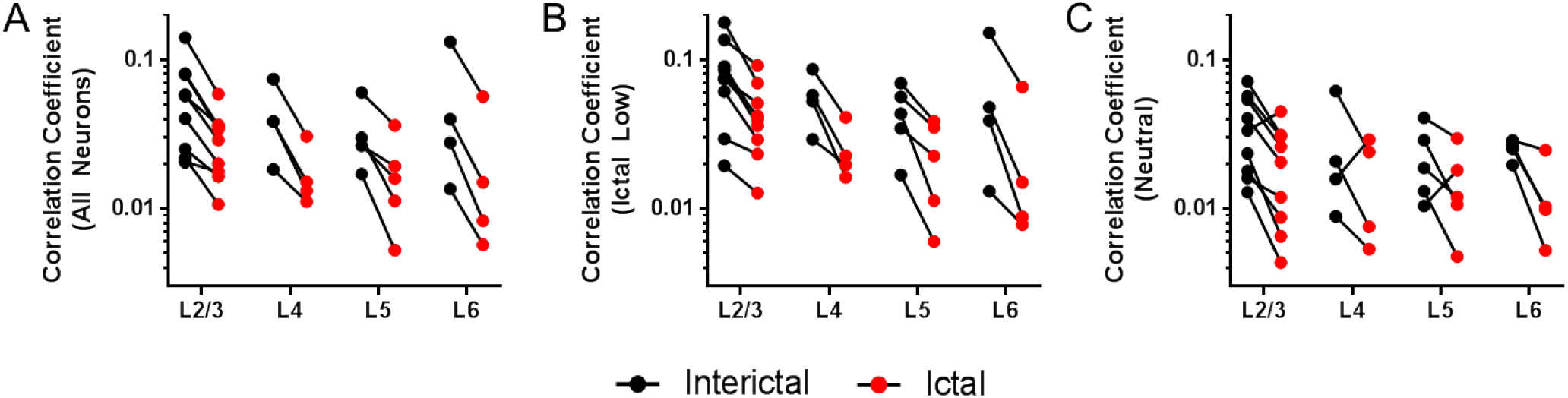
Deconvolution of the calcium signal yields similar correlations. Pairwise Pearson correlation coefficients using deconvolved data showed similar reduced correlations during the ictal state, with reductions in overall synchrony **(A)**, most pronounced in ictal-low neurons **(B)**, and a mixture of increased and decreased correlations with neutral neurons **(C)**.

**Supplementary Figure 8.**
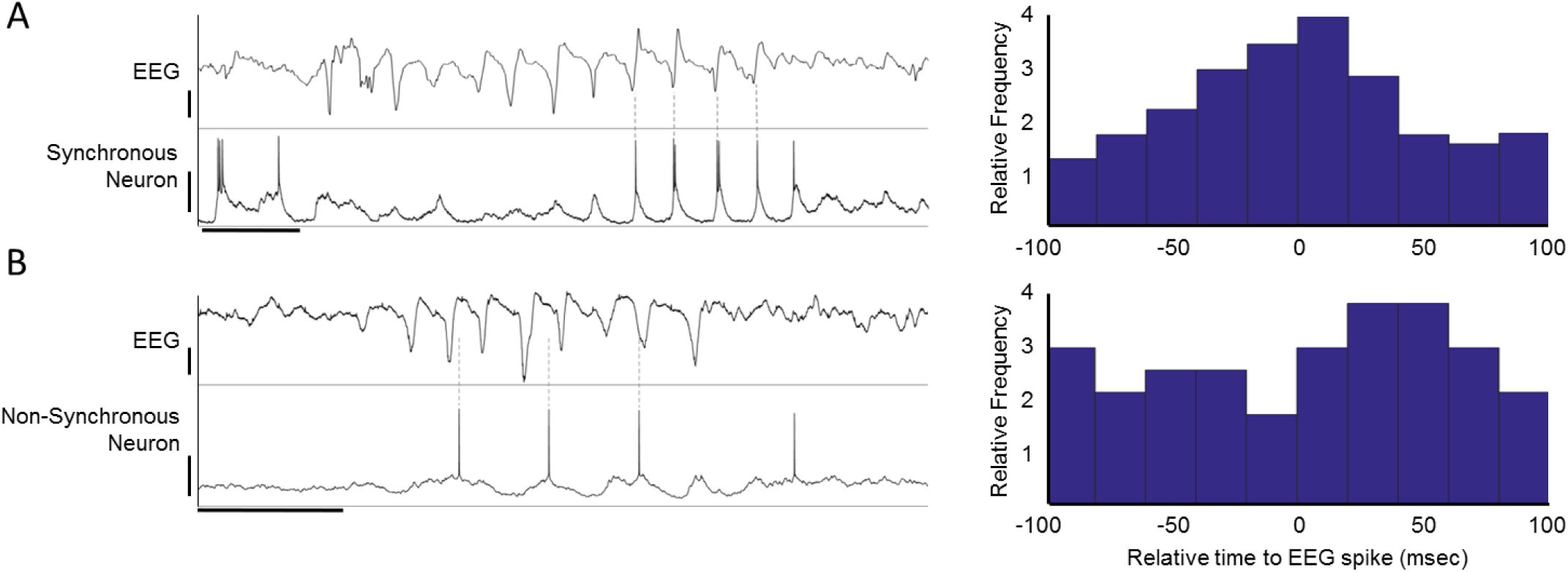
Patch-EEG synchrony. Layer 2/3 patched neurons had variable synchrony with EEG spikes. **(A)** Example of relatively high synchrony, with 29.3% of all EEG spikes coinciding with an action potential within ±20 msec. Note that several EEG spikes are not associated with an action potential. **(B)** Example of relatively low synchrony, with only 2.2% of all EEG spikes coinciding with an action potential within ±20 msec. Peri-spike time historgrams for each neuron are shown to the right. Time bar = 0.5 seconds; EEG amplitude bar = 20 µV; Patch amplitude bar = 40 mV. Overall, the likelihood of a neuron spiking within ±20 msec around an EEG spike was 17.9±3.9% (mean±SEM), indicating that synchrony of neurons during spike-wave seizures, measured at high temporal resolution, is low.

**Supplementary Figure 9.**
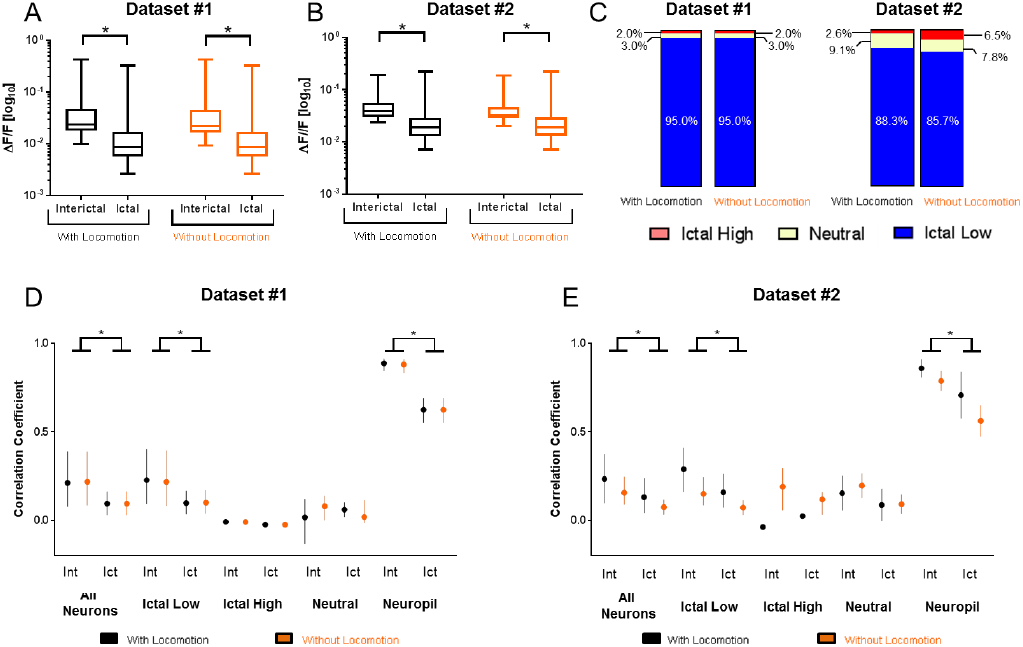
No significant change in neuron activity or synchrony when episodes of locomotion are removed (n = 2 datasets). To exclude possible movement contamination, when all imaging frames which occurred during wheel motion are removed, there is no significant change in: reduction in overall activity in neurons (n=100 and n=77 in Dataset #1 (A) and #2 (B), respectively; *p<0.0001, Kruskal-Wallis test with Dunn’s test for multiple comparisons), (C) relative proportion of ictal low neurons compared to ictal high and neutral neurons, and significant reduction in synchrony of all neurons, ictal low neurons and neuropil in the ictal state in Dataset #1 (D) and #2 (E), respectively; \**p*<0.0001, one-way ANOVA with Bonferroni test for multiple comparisons.

